# Transcriptomic responses of *Vibrio campbellii* to quorum sensing inhibition and antibiotic stress

**DOI:** 10.64898/2025.12.10.693564

**Authors:** Norfarrah Mohamed Alipiah, Natrah Fatin Mohd. Ikhsan, Hirzahida Mohd Padil, Sarmila Muthukrishnan

**Affiliations:** Universiti Putra Malaysia; University of Malaya: Universiti Malaya

**Keywords:** quorum sensing, quorum sensing inhibitor, RNA-seq, transcriptome, *Vibrio campbellii*, antimicrobial resistance

## Abstract

Quorum sensing (QS) is a key regulatory system in *Vibrio campbellii* that modulates virulence and communal behaviours. This study investigated the early transcriptional responses to a quorum sensing inhibitor (QSI) and tetracycline using RNA sequencing (RNA-seq). Following one hour of exposure, 130 and 539 differentially expressed genes (DEGs) were identified under QSI and tetracycline treatments, respectively. Quorum sensing inhibitory activities leading to upregulation of sulfur and porphyrin metabolism genes and downregulation of transporters and QS-related pathways, while tetracycline exposure induced genes related to ribosomal activity, DNA replication, and stress responses. KEGG and GO enrichment analyses revealed distinct pathway regulatory patterns, including in quorum sensing, two-component systems, and biofilm formation. These results highlight the mechanistic distinctions in bacterial response to quorum sensing interference versus antibiotic stress, emphasizing the potential of QSIs as precision anti-virulence agents that modulate pathogenic behaviour while minimizing the selective pressure associated with conventional antibiotics.

## Introduction

*Vibrio campbellii* is a gram-negative bacterium commonly found in marine environment and poses a significant threat to aquatic organisms due to its pathogenic nature (1). The pathogenesis of *V. campbellii* is mediated by quorum sensing (QS), a bacterial cell-to-cell communication mechanism mediated by the production and detection of small signalling molecules known as autoinducers (AIs). As bacterial cell density increases, the concentration of AIs also accumulates in the surrounding environment, leading to the activation of QS-controlled genes involved in virulence, motility and biofilm formation (2–4). Disruption of the QS through the utilization of quorum sensing inhibitors (QSIs) has emerged as a promising strategy for controlling bacterial infection by attenuating virulence without exerting strong selective pressure associated with antibiotics (5).

*Vibrio campbellii* utilizes a complex QS system involving multiple autoinducers, making it an ideal model to explore antivirulence strategies. Recent transcriptomic studies have shown that treatment with a specific QSI such as sodium ascorbate significantly downregulates genes associated with virulence and biofilm formation in *V. campbellii*, highlighting the potential of QSIs as therapeutic alternatives to conventional antibiotics (6)

Various QSIs have been identified from natural sources including marine algae and medicinal plants. These compounds disrupt QS by inhibiting signal biosynthesis or blocking receptor binding, thereby interfering coordination of pathogenic behaviours (7). In contrast, antibiotics such as tetracycline target essential cellular processes including protein synthesis that often triggered broad stress responses (8). The widespread and frequently indiscriminate use of antibiotics in aquaculture has contributed to the alarming rise of antimicrobial resistance (AMR), driving the search for alternative or complementary antimicrobial strategies. In parallel, molecular investigations into host responses to *Vibrio* infection have shown that shrimp haemocytes activate innate immune pathways, including pattern recognition receptors (PRRs), to detect and respond to bacterial invasion (9). These finding reinforce the importance of understanding both host and pathogen transcriptional responses in developing sustainable disease control strategies in aquaculture.

Advancement in transcriptomic technologies, particularly RNA sequencing (RNA-seq) now enable high-resolution profiling of bacterial responses to chemical stressors. Transcriptomic analysis identifies differentially express genes (DEGs) and reveals cellular pathways that are activated or repressed upon treatments. In the context of antibiotic exposure, previous studies has shown that bacteria often up-regulate stress response genes, including those related to efflux pump, DNA repair, and oxidative stress mitigation (10, 11). Similarly, QSIs have been shown to modulate distinct transcriptional network, including those involved in metabolism, membrane structure, and signal transduction (12, 13).

Although antibiotics remain central to bacterial control, the growing antimicrobial resistance (AMR) crisis necessitates for deeper molecular insight into bacterial adaptation. Comparative transcriptomic profiling response. to both QSIs and antibiotics could uncover key regulatory targets and guide the design of next-generation antimicrobials (14, 15). This study investigated the early transcriptional response of *V. campbellii* to QSI and tetracycline treatment using RNA-seq. Differentially express genes (DEGs) and enriched pathways across treatments were analyzed to elucidate the distinct cellular programs activated by QS disruption versus translational inhibition. These findings will contribute to our deeper understanding of pathogen regulation and support the development of effective antivirulence therapies.

## Materials and methods

### Bacterial strains and culture conditions

*Vibrio campbellii* BB120 was maintained in glycerol stock at −80 °C and revive on TCBS agar plates (Oxoid, UK) plates. A typical single colony was inoculated into 5 mL tryptic soy broth (TSB; Oxoid, UK) supplemented with 1.5% NaCl (w/v) and incubated at 30 °C with shaking at 200 rpm. The overnight culture was diluted 1:10 in fresh TSB with 1.5% NaCl and incubated until the culture reached mid-log phase (OD600 ∼ 0.5).

### Experimental treatments conditions

Three treatment condition were tested: (1) QSI extract from *Halamphora* sp. (acetone extract) at 0.2 mg/mL (16), (2) tetracycline at 3 µg/mL, and (3) phosphate-buffer saline (PBS; Invitrogen, USA) as control. The QSI extract and tetracycline were prepared by diluting stock solutions (2 mg/mL and 30 µg/mL, respectively) in PBS. For each treatments, 9 mL of mid-log phase culture was transferred into 50 mL falcon tubes and treated accordingly. Cultures were incubated at 30 °C with shaking at 200 rpm for 1 hour. Cells were harvested by centrifugation at 4,000 x g for 5 minutes. Pellets were resuspended in RNAlater (Invitrogen, USA) and stored at −80 °C until RNA extraction. A total of three biological replicates were prepared per treatment condition (QSI, tetracycline, and PBS control). Each biological replicate was sequenced in duplicate, generating 18 RNA-seq datasets for downstream analysis

### RNA extraction and quality

Total RNA was extracted using RNA extraction kit (Macherey-Nagel, Germany) following the manufacturer’s protocol. Briefly, cell pellet were lysed using lysis buffer and filtered using the NucleoSpin® Filter columns by centrifugation for 1 min at 11,000 x g. The flowthrough was precipitated with ethanol. The lysate were loaded onto NucleoSpin® RNA Column, centrifuge for 1 min at 11,000 x g. The RNA on binding column were proceeded with desalted and rDNase enzyme treatments followed by washing. RNA was eluted in 50 µL with RNase-free water. RNA quality and concentration were assessed using Jenway Genova spectrophotometer (Cole-Parmer, USA). Samples with a 260/230 ratio between 2.0-2.2, 260/230 ration between 2.0-2.2, and >200 ng/µL were selected for sequencing.

### Library preparation and RNA sequencing

RNA samples were shipped to NextGen Genomic Laboratory (Malaysia). RNA Integrity was confirmed using Agilent TapeStation (Agilent, USA) with RIN values >7. rRNA depletion was performed using NEBNext rRNA Depletion kit (Bacteria) and libraries were prepared using MGIEasy RNA Directional Prep Kit (MGI Tech, China). cDNA synthesis, adapter ligation, and PCR amplification were conducted following manufacturer’s protocol. Fragment size distribution (300 - 500 bp) was confirmed using D1000 ScreenTape, Sequencing was performed on the DNBSEQ-G400 platform using a High-throughput Rapid Sequencing Set (MGI Tech, China).

### RNA-seq data processing

Raw reads were assessed for quality using FastQC Version 0.11.8. Low quality reads (Phred score< Q20) and adapter sequences were removed using Cutadapt Version 3.50 (17) via Trim-galore Version 0.6.7. The resulting clean reads were then evaluated using MultiQC Version 1.14 (18). rRNA sequences were filtered out using SortMeRNA Version 4.3.6 (19). Cleaned reads were aligned to the *Vibrio campbellii* ATCC BAA-1116 reference genome (GenBank accession: GCA_000464435.1) using the STAR aligner Version 2.7. 10a (20), following genome indexing. Gene level quantification was performed with feature Counts Version 2.0.3 (21) using the gene annotation provided in GTF format from NCBI GenBank. Gene identifiers (e.g., *M892_XXXXX*) correspond to locus tags generated from the annotation file used in this study for the *V. campbellii* ATCC BAA-1116 genome.

### Differentially expressed gene analysis

Raw count data underwent quality checks at both the sample and gene levels. Principal Component Analysis (PCA) was used to assess sample clustering and detect outliers. Genes with low expression (mean counts < 10) were filtered out to improve statistical power. Differentially expression analysis was performed using DESeq2, which fits negative binomial distribution and applies Wald test for significance testing. Genes with an adjusted p-value < 0.05 and log2 fold change >1 or < 1 were considered significantly differentially expressed.

### Functional enrichment and pathway analysis

Gene Ontology (GO) and KEGG pathway enrichment analyses were performed using ClusterProfiler Version 4.2.2 (22). Both Over-Representation Analysis (ORA) and Gene Set Enrichment Analysis (GSEA) were applied. ORA identified pathway that overrepresented among DEGs, while GSEA assessed global expression trends without reliance on a DEG threshold. Enriched pathways were visualized using the Enrichplot package, presented in the form of gene concept networks and enrichment maps. Pathview Version 1.34 (23) was used to map DEGs onto KEGG pathways to aid biological interpretation.

## Results

### Transcriptome overview of *Vibrio campbellii* BB120 under QSI and tetracycline treatment

To investigate the early transcriptional response of *V. campbellii* BB120 to quorum sensing inhibition and antibiotic stress, mid-log phase cultures were exposed to either QSI extract (from *Halamphora* sp. acetone extract) or tetracycline for one hour.

RNA-seq generated a total of 162 million reads with high quality scores ( Q20 > 98%) and an average GC content of 47% across all samples (Table 1). Alignment statistics showed a mean length of aligned transcript 1,135 nucleotides, with a total read count 2.3 million mapped reads across all samples (Fig 1A). This indicates high-quality transcript coverage suitable for downstream analysis. Sequence data have been deposited in the Sequence Read Archive (SRA) under accession numbers: SRR31372477; SRR31372476; SRR31372475; SRR31372474; SRR31372473; SRR31372472; SRR31372471; SRR31372470; and SRR31372469.

**Fig 1.**
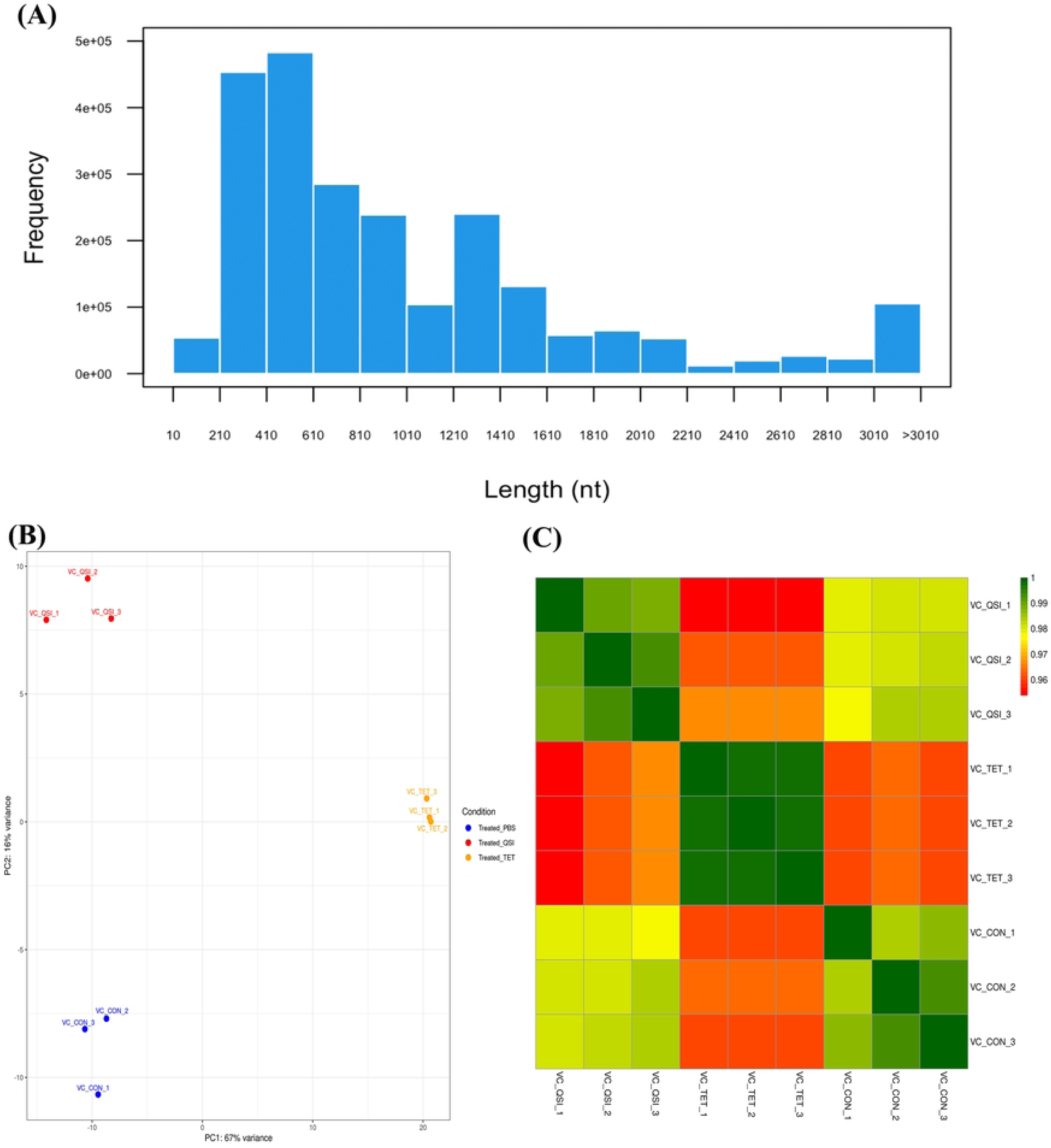
Analysis of aligned read counts, sample clustering, and correlation across treatments. In Fig 1A – 1C, group labels indicate the given treatments given to *V. campbellii* BB120. VC_CON (PBS control as treatment), VC_TET (tetracycline treatment) and VC_QSI (QSI extract treatment). (A) A histogram showing the frequency distribution of aligned reads count. (B) Principal component analysis (PCA) plot demonstrating that biological replicates cluster according to each treatment group. The in the axis labels indicate the percentage of principal component that explains the samples variance. (C) Correlation heatmap generated using the heatmap package, showing similarities among samples based on gene expression profiles.

**Table 1.**
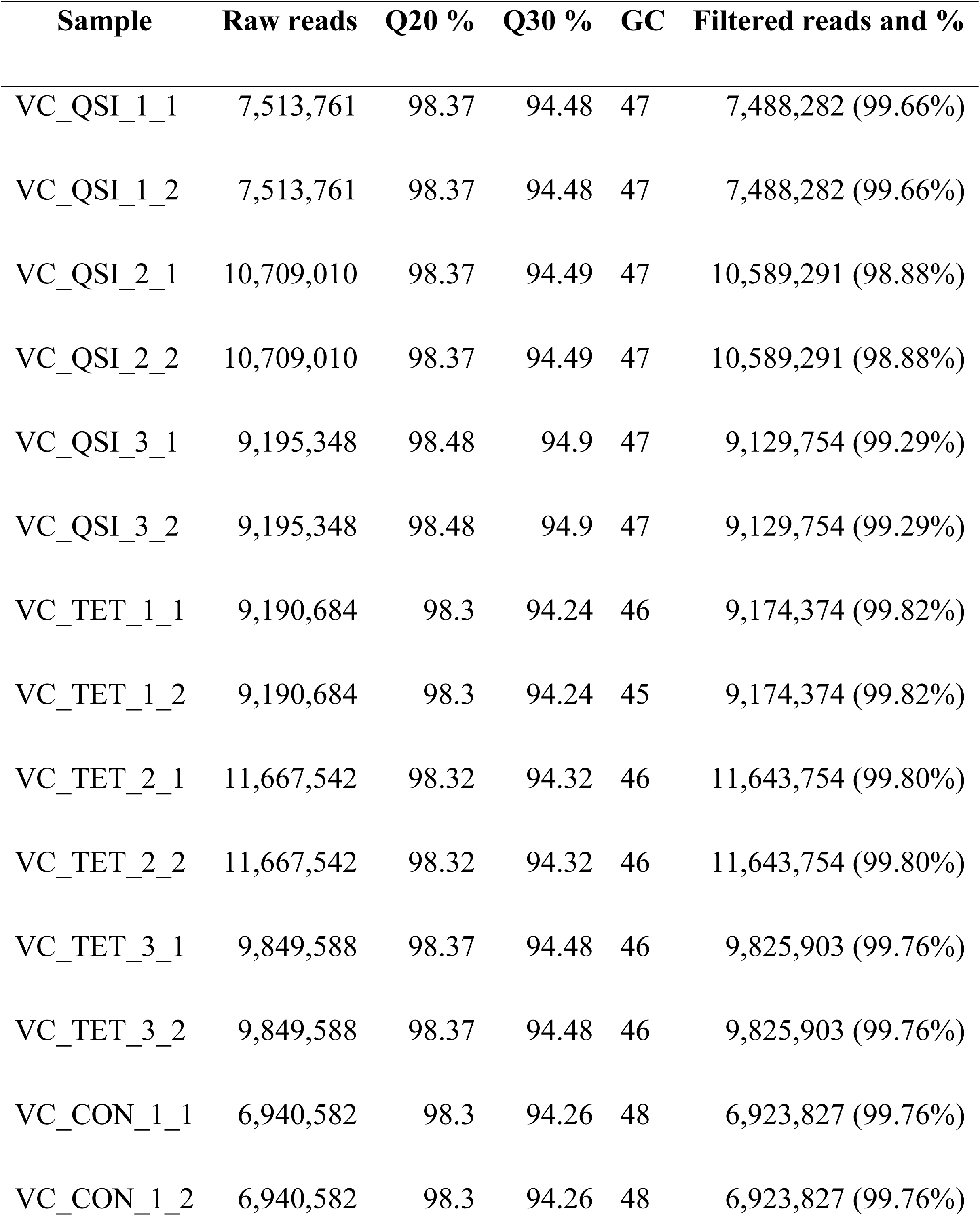
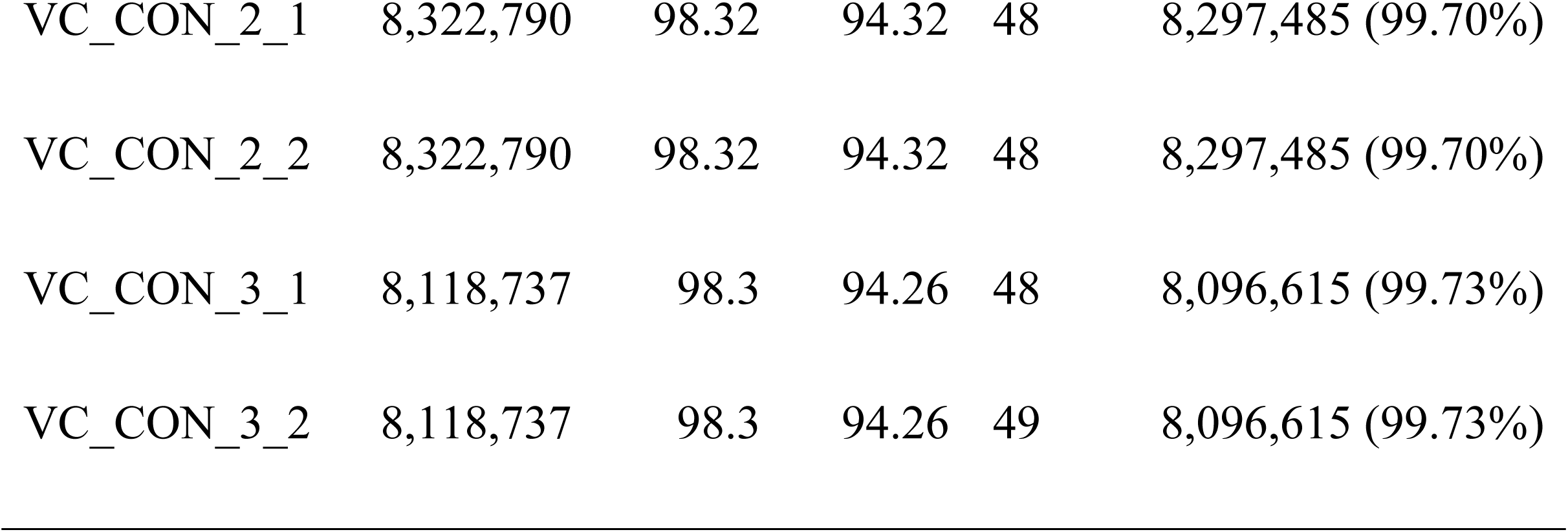
Data quality and filtering statistic for RNA-seq reads across all treatment.

| Sample | Raw reads | Q20 % | Q30 % | GC | Filtered reads and % |
| --- | --- | --- | --- | --- | --- |
| VC_QSI_1_1 | 7,513,761 | 98.37 | 94.48 | 47 | 7,488,282 (99.66%) |
| VC_QSI_1_2 | 7,513,761 | 98.37 | 94.48 | 47 | 7,488,282 (99.66%) |
| VC_QSI_2_1 | 10,709,010 | 98.37 | 94.49 | 47 | 10,589,291 (98.88%) |
| VC_QSI_2_2 | 10,709,010 | 98.37 | 94.49 | 47 | 10,589,291 (98.88%) |
| VC_QSI_3_1 | 9,195,348 | 98.48 | 94.9 | 47 | 9,129,754 (99.29%) |
| VC_QSI_3_2 | 9,195,348 | 98.48 | 94.9 | 47 | 9,129,754 (99.29%) |
| VC_TET_1_1 | 9,190,684 | 98.3 | 94.24 | 46 | 9,174,374 (99.82%) |
| VC_TET_1_2 | 9,190,684 | 98.3 | 94.24 | 45 | 9,174,374 (99.82%) |
| VC_TET_2_1 | 11,667,542 | 98.32 | 94.32 | 46 | 11,643,754 (99.80%) |
| VC_TET_2_2 | 11,667,542 | 98.32 | 94.32 | 46 | 11,643,754 (99.80%) |
| VC_TET_3_1 | 9,849,588 | 98.37 | 94.48 | 46 | 9,825,903 (99.76%) |
| VC_TET_3_2 | 9,849,588 | 98.37 | 94.48 | 46 | 9,825,903 (99.76%) |
| VC_CON_1_1 | 6,940,582 | 98.3 | 94.26 | 48 | 6,923,827 (99.76%) |
| VC_CON_1_2 | 6,940,582 | 98.3 | 94.26 | 48 | 6,923,827 (99.76%) |
| VC_CON_2_1 | 8,322,790 | 98.32 | 94.32 | 48 | 8,297,485 (99.70%) |
| VC_CON_2_2 | 8,322,790 | 98.32 | 94.32 | 48 | 8,297,485 (99.70%) |
| VC_CON_3_1 | 8,118,737 | 98.3 | 94.26 | 48 | 8,096,615 (99.73%) |
| VC_CON_3_2 | 8,118,737 | 98.3 | 94.26 | 49 | 8,096,615 (99.73%) |

Principal Component Analysis (PCA) revealed clear separation among QSI, tetracycline and PBS treated samples, accounting for 83% of the total variance across PC1 and PC2 (Fig 1B). Hierarchical clustering analysis further confirmed that biological replicates grouped according to treatment conditions (Fig 1C).

### Differential expression gene (DEG) profiles

Differential expression analysis identified 130 DEGs in QSI-treated samples and 539 DEGs in tetracycline-treated samples. each compared with the PBS control. The complete list of DEGs, including log2 fold change and adjusted p-value, is provided in S1 Table. In the QSI group, 70 genes were significantly upregulated and 60 downregulated, while in the tetracycline group 267 gene were upregulated and 271 downregulated.

These expression changes are visualized in MA plots (Fig 2A - B) and volcano plots (Fig 2C - D). Heatmap of the top 100 DEGs highlighted consistent expression pattern across biological replicates (Fig 3A - B).

**Fig 2.**
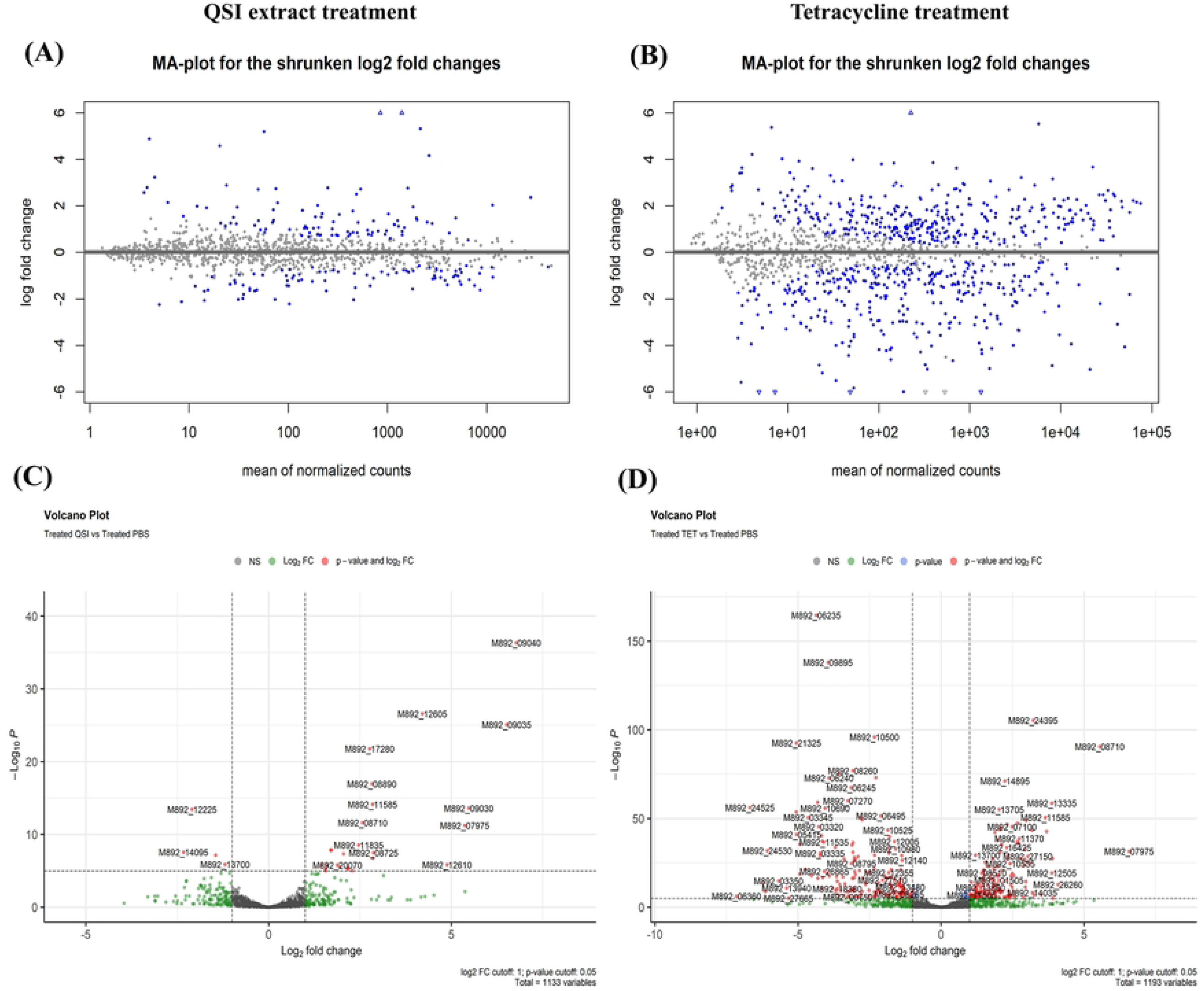
MA plot and Volcano plots of Differentially Expressed Genes (DEGs) pairwise comparison. (A) and (B) MA plots of DEGs distribution. The X axis represents the mean of normalized counts for DEGs and the Y axis represents the log fold change. Blue points represent genes with significant expression changes. (C) and (D) Volcano plot of DEGs significance and magnitude. The X axis represents Log_2_ fold change and the Y axis represents -Log_10_P p-value. Green and red points outside the vertical grey threshold bars represent genes with fold change >1 or <-1. Red points also represent up-regulated genes with adjusted p-value <0.05, highlighting the statistically significant DEGs.

**Fig 3.**
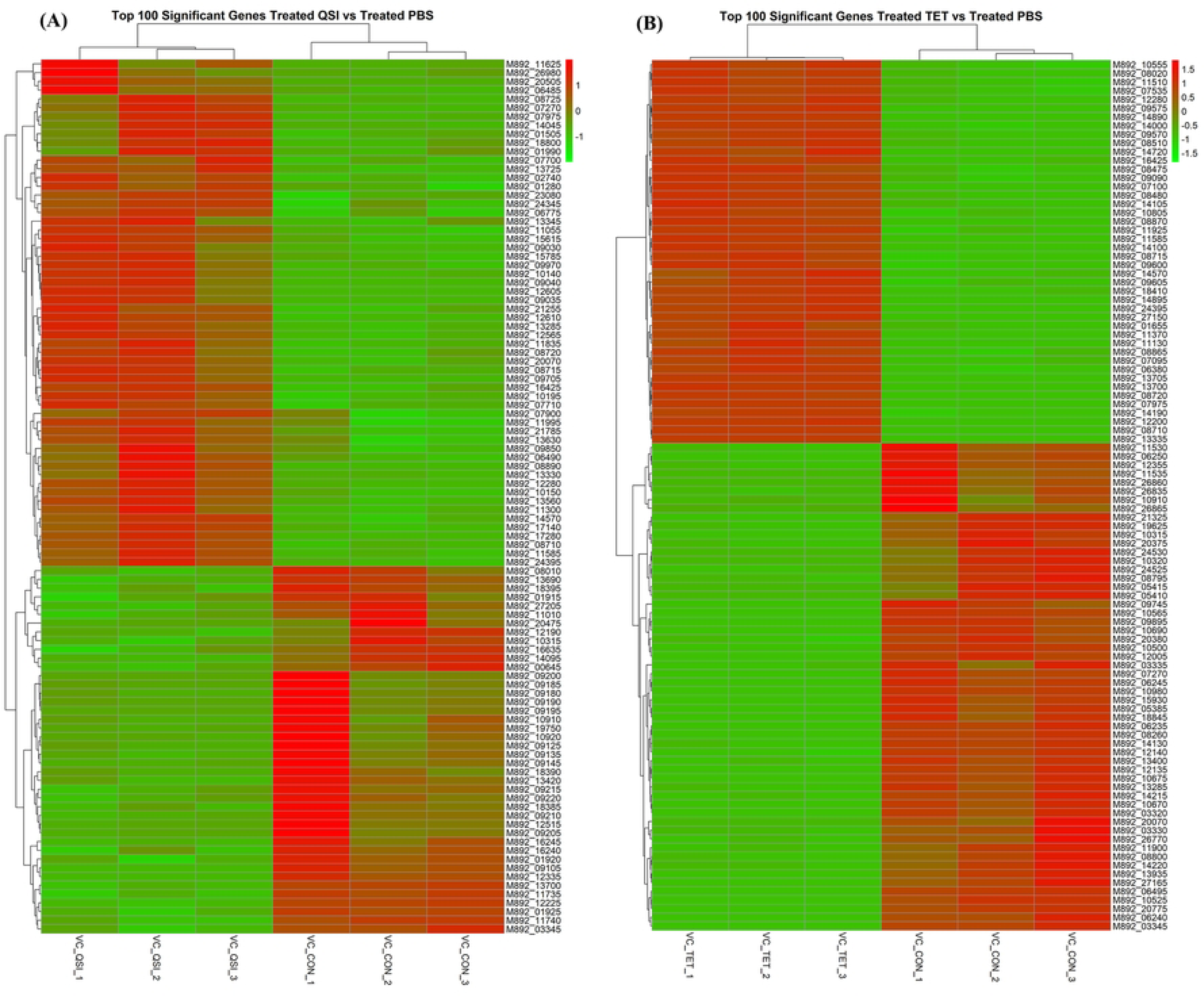
Transcriptional response of *V. campbellii* BB120 treated at mid-log phase for 1 h. (A) Clustered heatmap of top 100 significant DEGs under QSI extract treatment. (B) Clustered heatmap of top 100 significant DEGs under tetracycline treatment. Each column represent one sample, and each row represents one gene. The red and green gradients indicate up- and down-regulated expression, respectively.

The top ten upregulated and downregulated genes in each treatment are listed in Table 2. In the QSI group, strongly upregulated genes included *cyc*D, *cys*H, *cys*K, *cys*N, and *cys*I, while downregulated genes included transporter-associated genes *bam*B, *hyp*E, *exb*B, and *pot*B. In the tetracycline group, strongly upregulated gene included *mtl*A, *hol*D, *ubi*C, *tsa*D, *cyb*C, and *htp*X, whereas downregulated genes included *asn*B, *pha*C, *rpl*W, *kat*G, and *acs*.

**Table 2.**
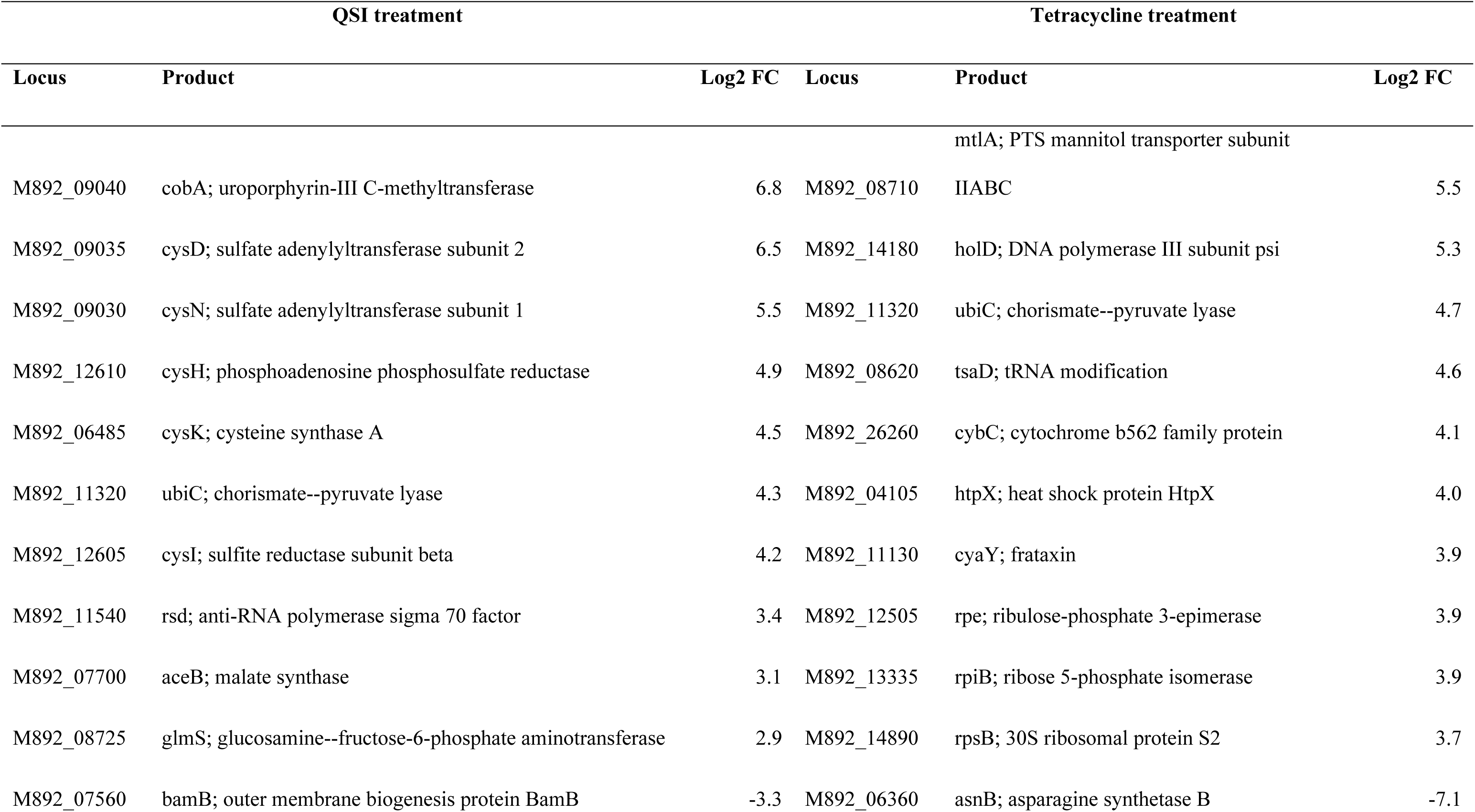
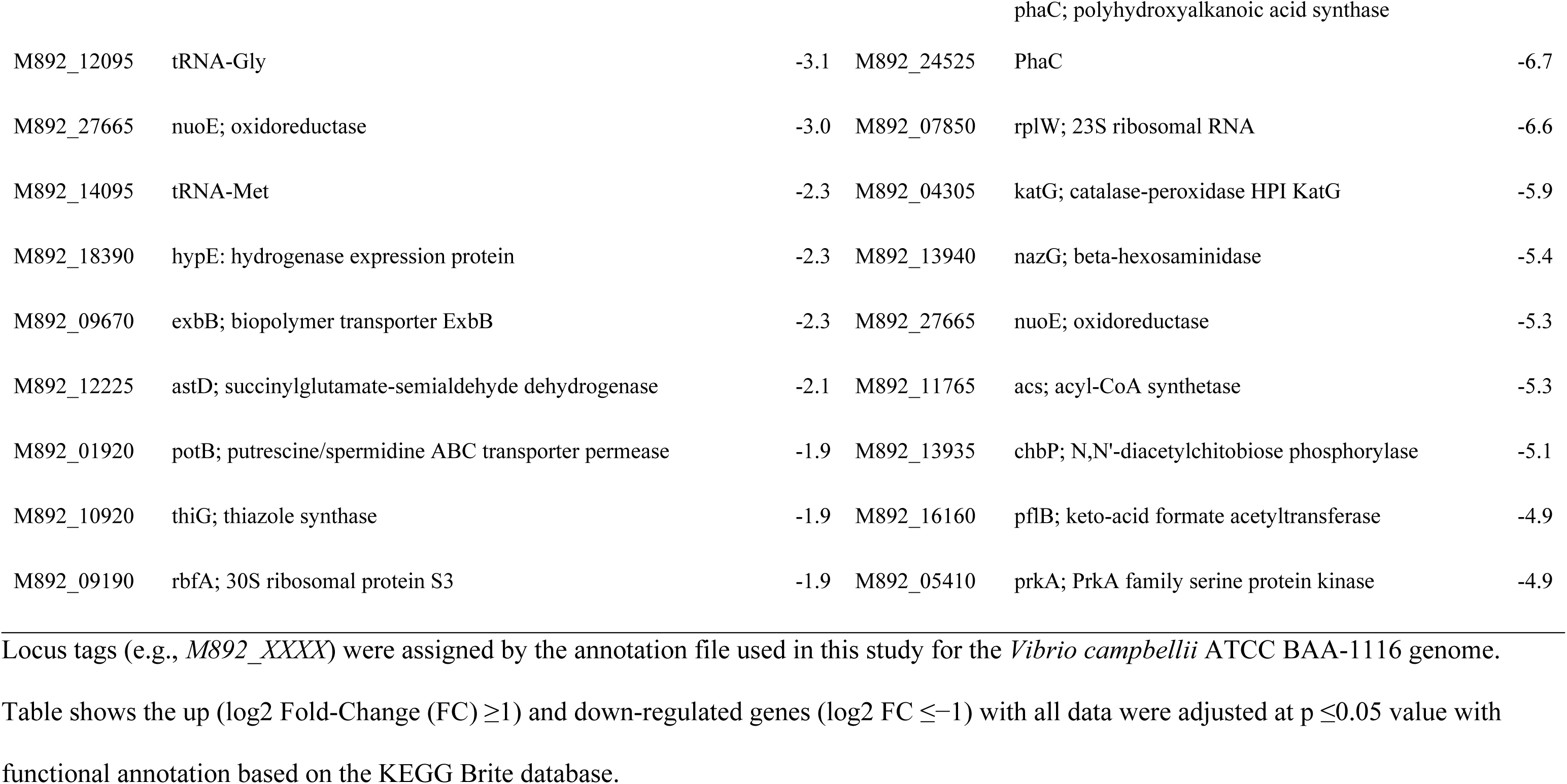
Top ten upregulated and downregulated genes in QSI and tetracycline treatments.

### Functional enrichment and gene network analysis

Gene Set Enrichment (GSEA) and Gene Concept Networks (GCNs) revealed distinct transcriptional responses of *V. campbellii* to QSI and tetracycline treatments (Fig 4a - d). Significantly enriched KEGG pathways identified through GSEA for each treatment are listed in S2 Table. In the QSI-treated group, pathways related to sulfur metabolism, ribosomal function, and microbial defense were significantly upregulated, while glutathione metabolism and phosphotransferase systems were significantly downregulated. In the tetracycline-treated group, enriched pathways included secondary metabolite biosynthesis, nitrogen metabolism, and amino acid biosynthesis, while butanoate and terpenoid metabolism were downregulated.

**Fig 4.**
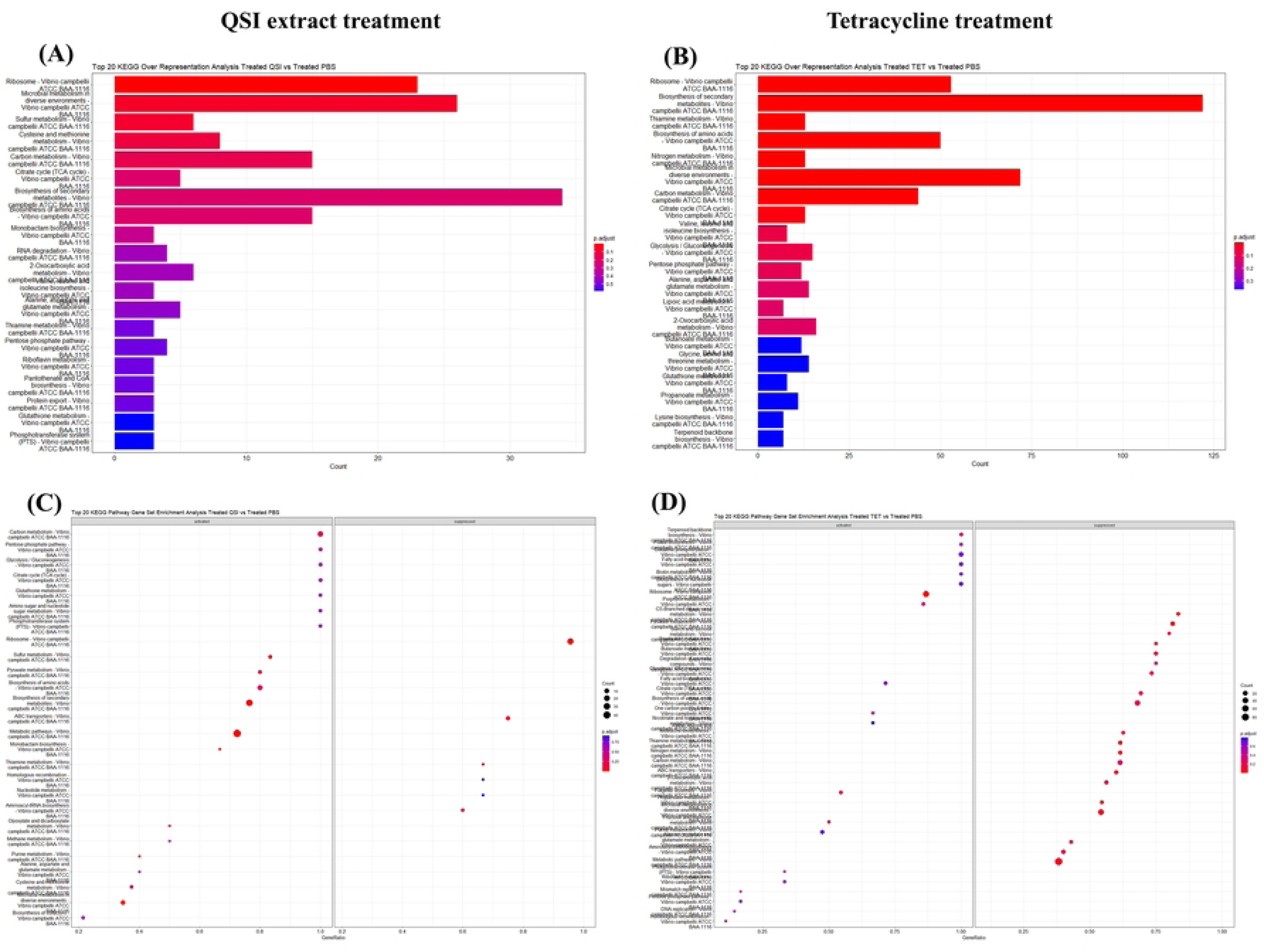
Gene set enrichment analysis (GSEA) and over representation analysis (ORA) for comparison condition based on KEGG terms. (A) and (C) ORA and GSEA of DEGs under QSI extract treatment. (B) and (D) ORA and GSEA of DEGs under tetracycline treatment.

GCN and enrichment maps (Fig 5A - D) showed treatment specific clustering of genes. Under QSI treatment, major co-expressed modules were associated with biofilm formation, ABC transporters, and amino sugar metabolism. In tetracycline-treated cells, enriched modules included butanoate metabolism, porphyrin pathways, and aminoacyl-tRNA biosynthesis.

**Fig 5.**
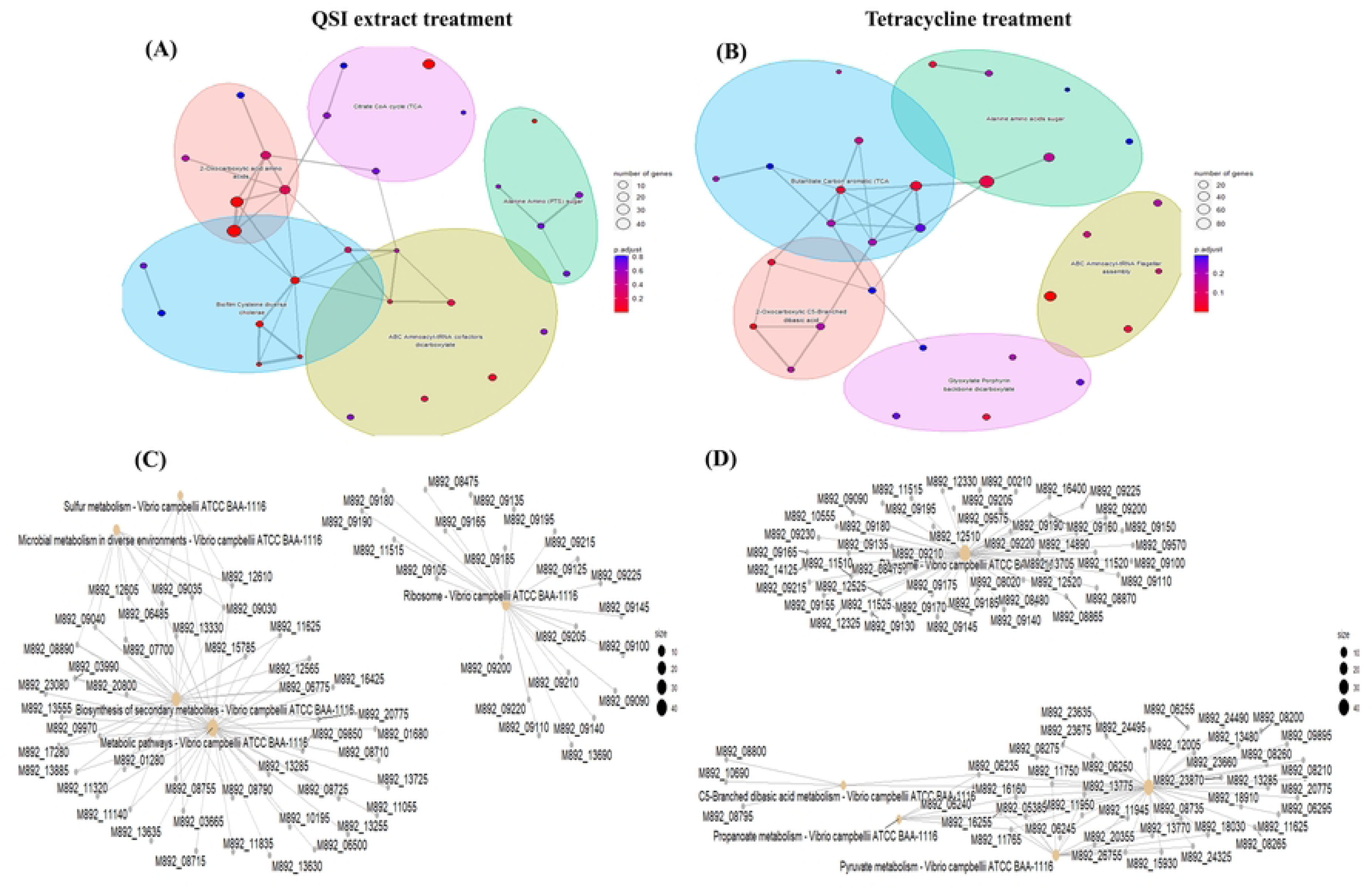
Network plots for gene set enrichment analysis. (A) and (C) Gene co-expression network under QSI extract treatment. The pathways included citrate cycle (TCA), ABC transporters, and sulfur metabolism indicate a shift in core energy metabolism, nutrient uptake, and redox balance in response to quorum sensing disruption. (B) and (D) Gene co-expression network under tetracycline treatment. In contrast to QSI treatment, pathways enhanced in this treatment are associated with glyoxylate metabolism, amino acid transport, and ribosome function, which leading to increased expression of stress response and survival-related genes under antibiotic pressure.

### KEGG pathway mapping and core regulators

KEGG pathway analysis revealed that both QSI and tetracycline treatments significantly altered core regulatory and metabolic process in *V. campbellii* (Fig 6 - 8). Key pathways affected two-component systems (TCS) (Fig 6), quorum sensing (QS) (Fig 7), and biofilm formation (Fig 8), in addition to sugar metabolism, the pentose phosphate pathway, and glycolysis/gluconeogenesis.

**Fig 6.**
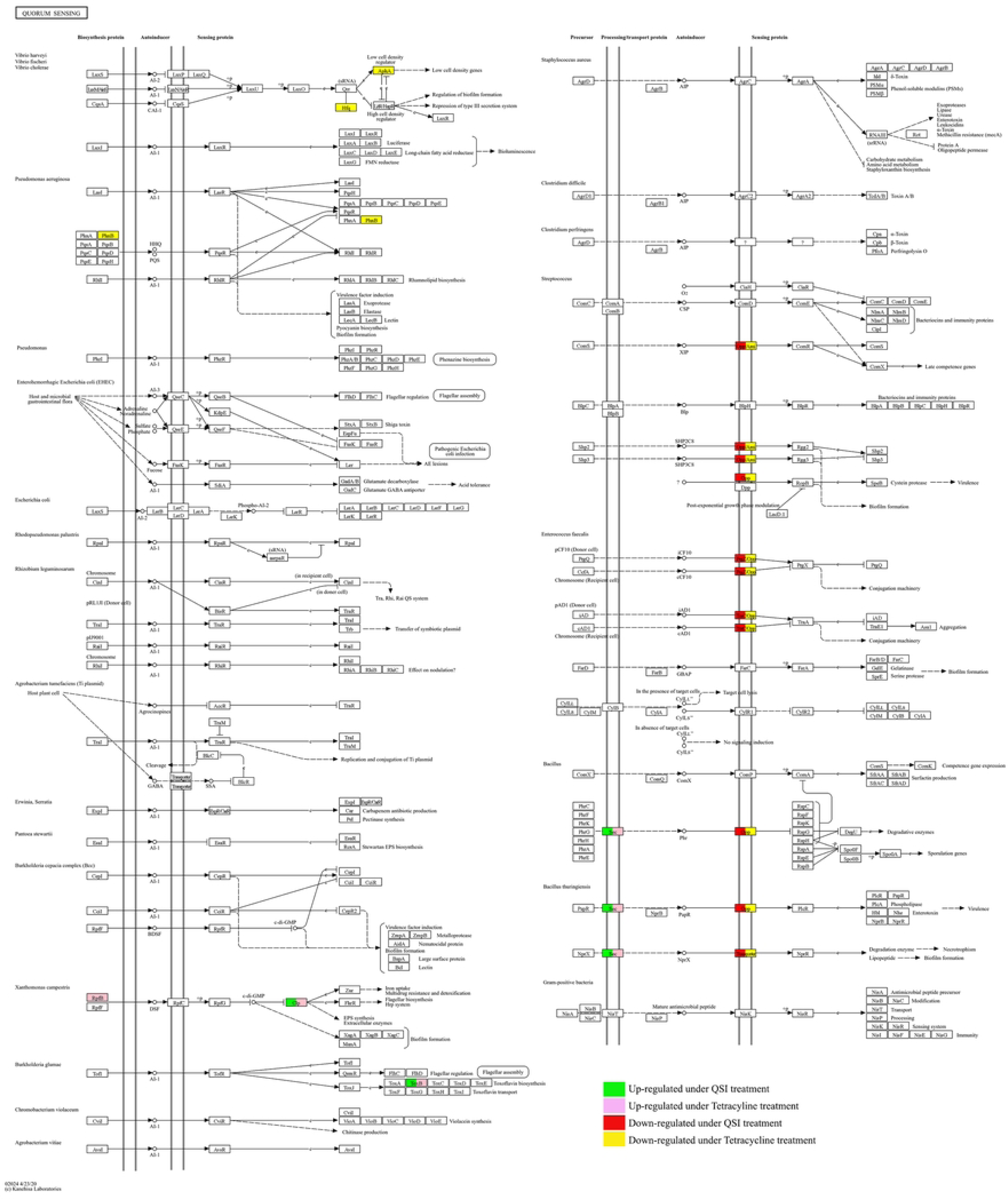
Enrichment of differently expressed genes (DEGs) in *Vibrio campbellii* across Quorum sensing pathway (vca02024) KEGG pathway. For QSI treatment, green boxes indicate that the genes are up-regulated, while red boxes shows down-regulation of the genes. For tetracycline treatment, the pink boxes indicate up-regulated genes and yellow boxes represents down-regulated genes.

**Fig 7.**
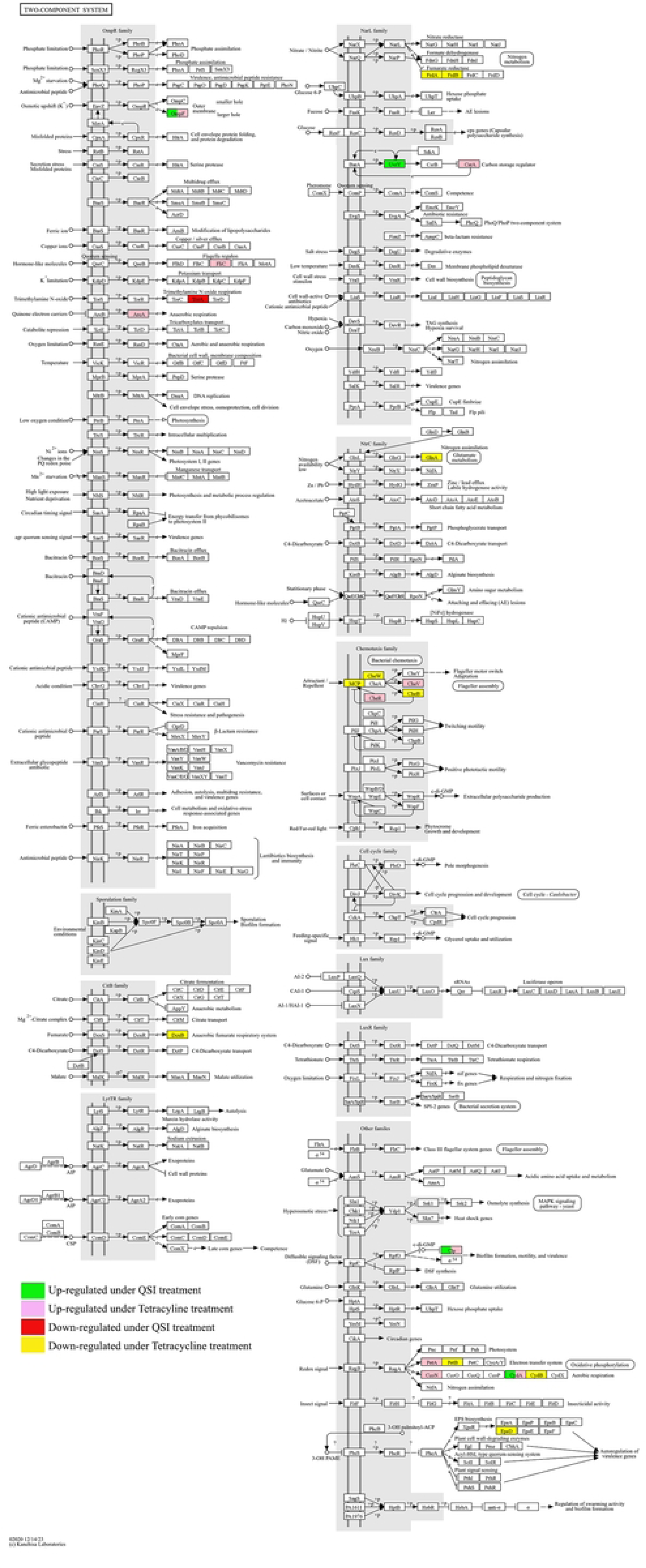
Enrichment of differently expressed genes (DEGs) in *Vibrio campbellii* across. Two-**component system pathway (vca02020) KEGG pathway**. For QSI treatment, green boxes indicate that the genes are up-regulated, while red boxes shows down-regulation of the genes. For tetracycline treatment, the pink boxes indicate up-regulated genes and yellow boxes represents down-regulated genes.

**Fig 8.**
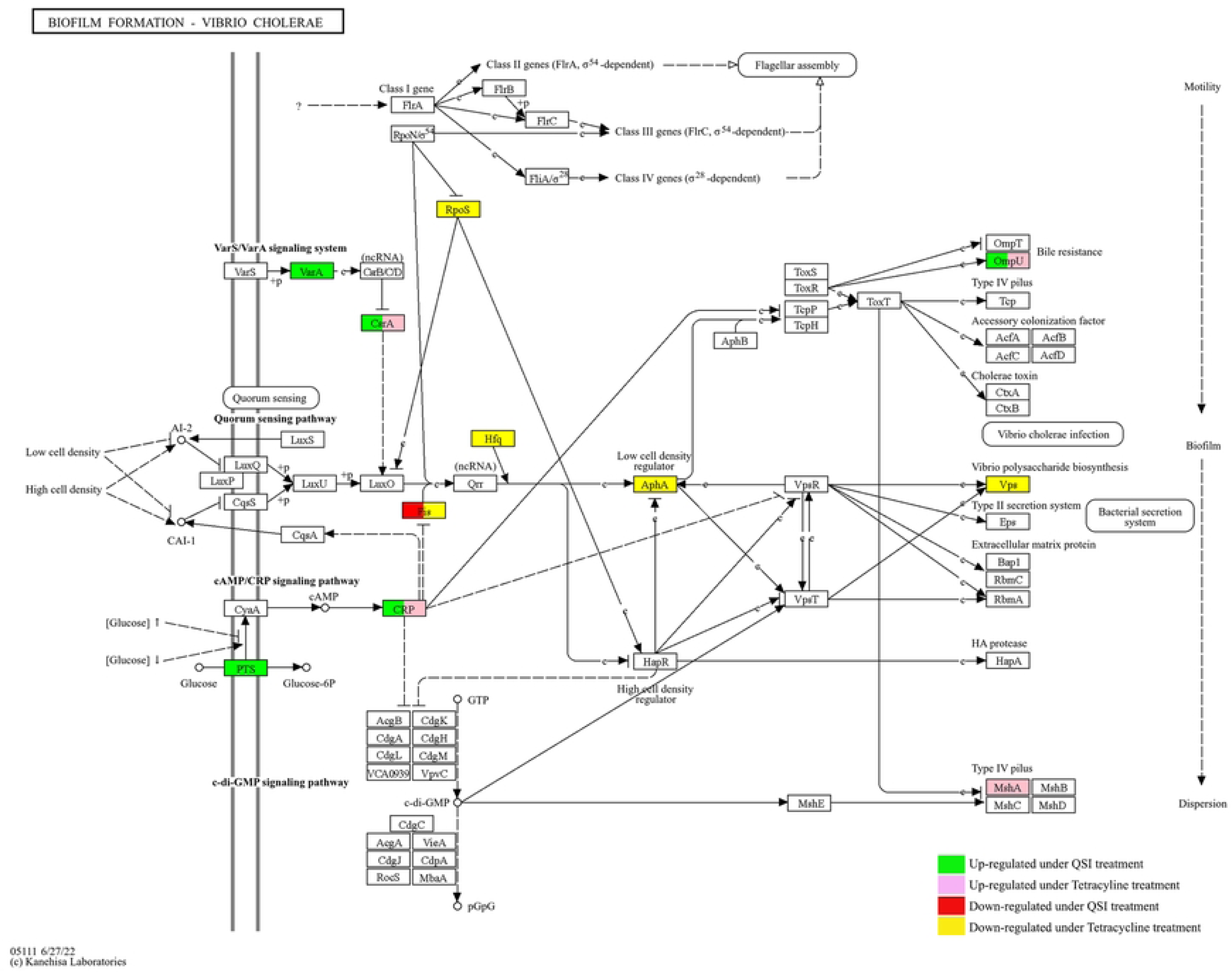
Enrichment of differently expressed genes (DEGs) in *Vibrio campbellii* across Biofilm formation (vca05111) KEGG pathway. For QSI treatment, green boxes indicate that the genes are up-regulated, while red boxes shows down-regulation of the genes. For tetracycline treatment, the pink boxes indicate up-regulated genes and yellow boxes represents down-regulated genes.

Within the TCS pathway, several regulatory genes were differentially expressed. Both treatments upregulated global regulators such as *csr*A, *ompr*F, *crp*, and *cyd*A, while tetracycline specifically induced motility and chemotaxis related genes (*fli*C, *che*R) and repressed genes linked to respiration (*dcu*B, *frd*A/B) and exopolysaccharide biosynthesis (*wec*C).

Transcriptomic changes were also observed in QS-associated genes. QSI treatment downregulated *opp*A and *mpp*A, upregulated *sec*DF whereas both treatment upregulated *crp* and *rib*A. Tetracycline uniquely induced genes related to fatty acid and tryptophan metabolism (*fad*F, *trp*G) as well as regulators *aph*A, and *hfq*.

Biofilm-related pathways were differentially modulated by treatments. QSI exposure upregulated genes involved in early biofilm development, including *sir*A, *csr*A, and *omp*U, while both treatments downregulated *fis*. In contrast, tetracycline broadly repressed genes linked to stress adaptation and matrix biosynthesis (*rpo*S, *hf*q, *aph*A, and *vps*U), but selectively inducer *msh*A, associated with initial surface attachment.

## Discussion

Aquaculture continue to face increasing challenges from bacterial pathogens including from *Vibrio* spp., which caused significant economic and ecological losses. Traditional reliance on antibiotics such as tetracycline has raised concerns due to the emergence of antimicrobial resistance (24), underscoring the urgent need for alternative strategies. Quorum sensing inhibitors (QSIs), particularly those derived from natural sources, have gains attention as antivirulence agents that disrupt bacterial communication and pathogenicity while exerting lower risk of resistance development (25–29). In this context, transcriptomic profiling provide a powerful approach to uncover the molecular basis of bacterial adaptation to both QS disruption and antibiotic stress. Our comparative transcriptomic analysis of *V. campbellii* BB120 exposed to QSI and tetracycline provide new insights by which bacteria modulate regulatory circuits, metabolic pathways, and stress responses under distinct antimicrobial pressures.

### Treatment driven transcriptomic variation

In this study, mid-log phase cultures of *V. campbellii* BB120 were exposed to either a quorum sensing inhibitor (QSI) extract derived from *Halamphora sp.* acetone extract (16) or the antibiotic tetracycline for a short period to capture immediate transcriptional changes. The one-hour exposure window was chosen to reflect early adaptive responses, a strategy that has been applied in previous transcriptomic studies to uncover rapid stress-induced regulatory shift (13, 30). Tetracycline was selected as it remains one of the most widely used antibiotic in aquaculture and is known to impose strong selective pressure leading to antimicrobial resistance (24). In contrast, QSIs, many of which are derived from natural sources, are increasingly explored as antivirulence agents capable of disrupting bacterial communication with lower risk of resistance development (25–29).

The RNA-seq dataset generated in this study were of high quality, with over 162 million reads obtained, high Q20 scores (>98%), and consistent GC content, providing reliable coverage for downstream transcriptomic analysis. Principal Component Analysis (PCA) and hierarchical clustering further confirmed that treatment type was the dominant driver of transcriptomic variation, with clear separation between QSI, tetracycline, and control groups. This distinct clustering indicates that *V. campbellii* rapidly reprograms its transcriptome in response to both communication disruption and antibiotic stress. Similar treatment-driven transcriptomic shifts have been reported in studies examining bacterial response to QSIs (31, 32), antibiotics (14) and untreated or baseline conditions (33, 34). Collectively these results validate the robustness of the experimental design and reinforce that the transcriptional reprogramming observed reflects early adaptive responses to distinct stressors.

### Targeted transcriptomic response to QSI

The transcriptional response of *V. campbellii* BB120 to QSI exposure was relatively focused, with 130 DEGs identified compared to the control. Notably, genes associated with sulfur metabolism, including *cys*D, *cys*H, *cys*K, and *cys*I, were strongly upregulated. Sulfur assimilation is a key target of quorum sensing (QS)-linked regulatory networks that enable bacteria to adapt to environmental changes, and the induction of these genes suggest that QS disruption may trigger compensatory metabolic reprogramming (35, 36). In addition, upregulation of cobA, which is involved in porphyrin metabolism, points to potential cross-talk between sulfur and porphyrin pathways in response to QS interference (37).

In contrast, QSI treatment suppressed genes encoding several transporter components, including *bam*B, *hyp*E, *exb*B, and *pot*B. These transporters play critical roles in substrate translocation, energy coupling, and solute exchange under stress conditions (38–41). Their downregulation suggests that QSI exposure impairs nutrient uptake and stress adaptability. This observation is consistent with the findings of Zhang, Li (42), who reported similar repression of transport-related genes, including sulfate/thiosulfate and carbohydrate transporters, in *E. coli* O157 exposed to a QSI compound. Together, these results suggest that QS interference selectively alters metabolic and transporter pathways, weakening cellular adaptability without inducing a broad physiological shutdown.

### Broad transcriptomic reprogramming under tetracycline stress

In contrast to the targeted effects of QSI, tetracycline treatment induced a much broader transcriptional reprogramming in *V. campbellii*, with 539 DEGs identified, consistent with its global inhibitory effects on protein synthesis. Many of the most upregulated genes were linked to carbohydrate transport and metabolism, DNA replication, and stress adaptation. For example, *mtl*A, encoding the PTS mannitol transporter, was strongly induced, consistent with enhanced sugar uptake as compensatory energy strategy under stress (43). Similarly, upregulation of *hol*D (DNA polymerase III subunit psi) and *ubi*C (chorismate-pyruvate lyse)reflects increased demands for DNA synthesis and energy metabolism. Stress response genes such as *tsa*D (tRNA modification), *cyb*C (oxidative stress), and *htp*X (membrane protein quality control) were also elevated, suggesting coordinated cellular effort to stabilize protein translation and maintain envelope integrity (44).

Simultaneously, tetracycline significantly downregulated gene associated with amino acid biosynthesis, respiration, and ribosomal function. Strong repression of *asn*B (asparagine synthase B), *pha*C (PHB synthase), and *rpl*W (23S rRNA) indicates reduced biosynthetic activity as a means of conserving energy during translational arrest (45). Downregulation of *kat*G (catalase-peroxidase) suggest compromised oxidative stress defences, while suppression of *acs*, *pfl*B, and *nuo*E points to reduced aerobic and anaerobic respiratory capacity. These transcriptional shifts are consistent with stress responses that prioritize energy efficiency and redox balance in hostile conditions (46). Overall, tetracycline imposed a global reorganization of cellular functions, enhancing pathways that support survival while repressing energy-intensive process. Such a pattern reflects the classic bacterial survival strategy under translational-inhibiting antibiotics and highlights gene targets potentially linked to adaptive or persistence.

### Distinct pathway level responses to QSI and tetracycline

The enrichment analyses revealed that QSI and tetracycline triggered distinct pathway-level adaptations *in V. campbellii*. In the QSI-treated group, sulfur metabolism, ribosomal function, and microbial defence pathways were significantly upregulated, suggesting a targeted enhancement of metabolic adaptation and translational capacity in response to quorum sensing disruption. At the same time, glutathione metabolism and phosphotransferase systems were suppressed, indicating altered redox balance and reduced nutrient uptake. Such patterns are characteristic of bacterial stress responses inducing quorum sensing interference (35, 36).

In contrast, tetracycline treatment enriched pathways linked to secondary metabolite biosynthesis, nitrogen metabolism, and amino acid biosynthesis, while repressing butanoate and terpenoid metabolism. These changes reflect a transcriptional shift away from growth-oriented process toward survival-focused responses under translational stress, consistent with the mode of action of protein synthesis inhibitor (43, 44)

Gene Concept Networks (GCNs) further illustrated these divergent responses. Under QSI exposure, key co-expression modules were associated with biofilm formation, ABC transporters, and amino sugar metabolism, emphasizing regulatory networks that support surface attachment and environmental adaptation. By contrast, tetracycline-treated cells showed enrichment in modules related to butanoate metabolism, porphyrin pathways, and aminoacyl-tRNA biosynthesis. These changes suggest disrupted energy metabolism and impaired translational machinery, hallmarks of bacterial adaptation to antibiotic stress (45, 46)

### Regulatory and functional pathway adaptations under QSI and tetracycline stress

#### Overview of core pathway modulation

KEGG pathway mapping revealed that both quorum sensing inhibition (QSI) and tetracycline exposure profoundly altered the expression of core biological process in *V. campbellii*. The most affected pathways included quorum sensing (QS), two-component systems (TCS), and biofilm formation, alongside sugar metabolism, the pentose phosphate pathway, and glycolysis/gluconeogenesis. Notably, the QS and TCS pathways are closely interconnected, as QS signalling often involves TCS-mediated regulation of downstream genes associated with virulence and biofilm development (47, 48). The transcriptome responses observed in this study highlight the extensive reprogramming of signalling networks and adaptive behaviour in *V. campbellii* when challenged with either communication disruption or antibiotic pressure.

### Two-component system (TCS) responses

The TCS plays a pivotal role in bacterial adaptation by sensing environmental stimuli and coordinating appropriate regulatory responses that govern virulence, motility, and metabolic activity (49–51). Both QSI and tetracycline treatments triggered extensive transcriptional changes within system, although the patterns of regulation differed by treatments.

In both groups, core regulatory genes involved in nutrient sensing and stress adaptation – such as *csr*A, *omp*F, *crp*, and *cyd*A that were consistently upregulated. *csr*A, a global post-transcriptional regulator, modulates metabolism, motility, QS, and virulence, and its upregulation suggests broad stress-responsive reprogramming, consistent with reports in *Yersinia psedotuberculosis* and *E. coli* (52–54). Similarly, *omp*F, encoding a porin associated with osmotic regulation and antibiotic resistance, was induced under both treatments. Upregulation of this gene has been linked to ciprofloxacin resistance in *Salmonella enteriditis* treated with cinnamon oil (55).

Upregulation of crp (a cAMP receptor protein) and cydA (a component of cytochrome bd oxidase) further indicated the activation of stress-adaptive respiratory control. cydA upregulation has been previously reported under respiratory stress in Mycobacterium smegmatis (56, 57). Interestingly, while cydA increased under both treatments, cydB was selectively downregulated, indicating treatment-specific respiratory adjustment.

Tetracycline produced broader TCS-related transcriptional shifts consistent with inhibitory mechanism on protein synthesis (58). Notably, genes related to motility and chemotaxis (*fli*C, *che*R) were induced, potentially supporting motility-based escape from antibiotic stress (44). Conversely, genes involved in respiration (*dcu*B, *frd*A/B), nitrogen assimilation (*gln*A), and exopolysaccharides production (*wec*C) were repressed, reflecting a metabolic transition from growth to survival through resource reallocation (45, 46). Together, these findings demonstrate that *V. campbellii* activates overlapping yet distinct TCS-mediated stress responses under QSI and tetracycline pressure, where QSI eliciting target modulation and signalling pathways, and tetracycline inducing global reprogramming associated with translational arrest.

### Quorum sensing pathway modulation

The transcriptome data also revealed treatment-specific regulation within QS-related genes, highlighting divergent adaptive mechanisms to communication disruption versus antibiotic stress. Under QSI treatment, *opp*A and *mpp*A, encoding oligopeptide transport proteins, were downregulated. These transporters form part of the Opp system, which mediates peptide import and contributes to virulence and competences in QS-regulated bacteria (59, 60). Their repression suggests diminished intercellular signalling and nutrient exchange under QSI exposure. In contrast, *opp*A and *mpp*A were upregulated under tetracycline treatment, likely reflecting a compensatory mechanism to enhance nutrient uptake during stress.

QSI treatment also induced secDF, a component of the Sec-dependent protein translocation machinery, which support envelope integrity under stress (61). Similarly, *rib*A (riboflavin biosynthesis) and *crp* were upregulated under both treatments. The crp gene functions as a global regulator integrating environmental and metabolic cues with QS-mediated gene expression and virulence regulation (62, 63). This shared upregulation as a common adaptive mechanism, even when challenged by distinct stressors.

Tetracycline uniquely induced *fad*D (fatty acid metabolism) and *trp*G (tryptophan biosynthesis), both associated with metabolic remodelling during oxidative or antibiotic stress (64, 65). Moreover, induction of *aph*A, a QS repressor favouring individual survival at low cell density (66), and *hfq*, an RNA chaperone essential for stress response and QS regulation (67, 68), indicates a transcriptional shift toward survival and stress tolerance. Collectively, these patterns reveal that QSI provokes selective tuning of QS and transport pathways, whereas tetracycline drives global transcriptional suppression and activation of stress related genes.

### Biofilm formation and regulatory dynamics

Biofilm-associated genes displayed contrasting expression profiles under QSI and tetracycline exposure, underscoring different adaptive trajectories. Under QSI treatment, several gene promoting biofilm formation – *sir*A, *csr*A, the PTS EIIA component, *crp*, and *om*pU were upregulated. *sir*A acts as a response regulator promoting biofilm development through *csr*A regulation, which controls motility, surface attachment, and carbon metabolism (69–71). The induction of PTS system components suggests enhanced carbohydrate uptake supporting matrix biosynthesis (72). These results imply that, despite disrupting QSI, QSI may simultaneously trigger compensatory mechanisms reinforcing early biofilm establishment.

Both treatments upregulated *crp* and *omp*U indicating activation of conserved stress responsive pathways. *crp* integrates metabolic and QS signalling (62, 63), while *omp*U contributes to environmental sensing and surface attachment (73). Conversely, *fis*, which regulates DNA topology and biofilm dispersal during rapid growth, was downregulated under both treatments, suggesting a shift toward a sessile, low growth state (74).

Tetracycline exposure, however, caused more extensive suppression of key regulatory and biofilm-associated genes including *rpo*S, *hfq*, *aph*A, and *vps*U. *rpo*S encodes the stationary phase sigma factor essential for stress adaptation and biofilm maturation (75), while *hfq* and *aph*A mediate post-transcriptional and QS regulatory control (66, 68). Downregulation of *vps*U, critical for exopolysaccharides biosynthesis, indicates impaired biofilm matrix formation (76). Interestingly, *msh*A, encoding the MHSA pilin for initial surface attachment, was upregulated only in tetracycline-treated cells, suggesting a compensatory attempt to initiate biofilm formation despite inhibit matrix development.

Overall, QSI treatment maintained metabolic activity and promoted early biofilm-related regulatory shift, while tetracycline broadly suppressed biofilm maturation and regulatory flexibility. These contrasting outcomes illustrated how QS disruption primarily modulates communication and attachment without triggering cellular shutdown, whereas antibiotic impose systemic transcriptional repression. This distinction highlights the therapeutic promise of QSIs as antivirulence agents capable of attenuating pathogenic traits with reduced selection pressure for resistance

## Conclusion

This study demonstrates that *Vibrio campbellii* BB120 exhibits distinct transcriptional responses to quorum sensing inhibition and tetracycline stress. QSI exposure triggered targeted reprogramming of sulfur metabolism, transport system, and biofilm-associated pathways, while tetracycline induced a broad survival-oriented response characterized by enhanced energy acquisition, DNA repair, and suppression of biosynthetic and respiratory processes. Pathway-level analyses revealed that both treatments engaged two-component systems and global regulators, but with divergent effects on quorum sensing and biofilm development. These findings highlight the potential of QSIs as antivirulence agents that modulate regulatory networks without imposing the extensive metabolic disruption caused by antibiotics. While additional validation and physiological assays are needed, this work provides a valuable transcriptomic framework for understanding bacterial adaptation to communication disruption versus antibiotic pressure, and supports the development of alternative strategies to mitigate antimicrobial resistance in aquaculture.

## Acknowledgements

Generative AI tools were used for language editing only. Scientific content, analysis, and interpretation were performed by the authors. Ikhsan Natrah is also affiliated with the Microalgae-Biota Technology and Innovation Research Group (ALBIC), Universiti Putra Malaysia. Further information about the research group is available at albic.my.

## Author contributions

Conceptualization: NMA, HMP, SM. Investigation: NMA. Methodology: NMA. Visualization: NMA. Writing -original draft preparation: NMA, IN, HMP, SM. Writing - review and editing: NMA, IN. Supervision: NMA, IN. Funding: IN

## Ethical statement

This study did not involve any experiments on human participants or animals. Ethical approval was therefore not required

## Data availability

All RNA sequencing data generated in this study have been deposited in the NCBI Sequence Read Archive (SRA) under BioProject accession number PRJNA1187262. Individual SRA runs are available under accession numbers SRR31372469 - SRR31372477. All processed data supporting the findings of this study are included in the Supplementary Tables S1-S2.

## Supporting documents

**S1 Table. Full list of differential expressed genes.** Complete list of differential expressed genes (DEGs) identified in *Vibrio campbellii* BB120 under QSI and tetracycline treatments compared with the PBS control. The table includes gene identifiers, log2 fold changes, and adjusted p-value.

**S2 Table. Enriched functional pathways.** Significantly enriched KEGG and GO pathway identified through over-representation analysis (ORA) and gene set enrichment analysis (GSEA) for QSI and tetracycline treated *Vibrio campbellii.* The table lists pathways IDs, pathway name, enrichment scores, and associated genes

## References

1. Srisangthong I, Sangseedum C, Chaichanit N, Surachat K, Suanyuk N, Mittraparp-Arthorn P. Characterization and genome analysis of Vibrio campbellii lytic bacteriophage OPA17. Microbiology spectrum. 2023;11(2):e01623–22.

2. Waters CM, Bassler BL. Quorum sensing: cell-to-cell communication in bacteria. Annu Rev Cell Dev Biol. 2005;21:319–46.

3. Bridges AA, Bassler BL. The intragenus and interspecies quorum-sensing autoinducers exert distinct control over Vibrio cholerae biofilm formation and dispersal. PLoS biology. 2019;17(11):e3000429.

4. Simpson CA, Petersen BD, Haas NW, Geyman LJ, Lee AH, Podicheti R, et al. The quorum-sensing systems of Vibrio campbellii DS40M4 and BB120 are genetically and functionally distinct. Environmental microbiology. 2021;23(9):5412–32.

5. Vashistha A, Sharma N, Nanaji Y, Kumar D, Singh G, Barnwal RP, et al. Quorum sensing inhibitors as Therapeutics: Bacterial biofilm inhibition. Bioorganic Chemistry. 2023:106551.

6. Han B, Zheng X, Baruah K, Bossier P. Sodium ascorbate as a quorum-sensing inhibitor leads to decreased virulence in Vibrio campbellii. Frontiers in Microbiology. 2020;11:520947.

7. Tripathi S, Purchase D, Govarthanan M, Chandra R, Yadav S. Regulatory and innovative mechanisms of bacterial quorum sensing–mediated pathogenicity: a review. Environmental Monitoring and Assessment. 2023;195(1):75.

8. Chopra I, Roberts M. Tetracycline antibiotics: mode of action, applications, molecular biology, and epidemiology of bacterial resistance. Microbiology and molecular biology reviews. 2001;65(2):232–60.

9. Alipiah NM, Salleh A, Sarizan NM, Ikhsan N. Molecular characterization and gene expression of pattern recognition receptors in brown-marbled grouper (Epinephelus fuscoguttatus) fingerlings responding to vibriosis infection. Developmental & Comparative Immunology. 2024;161:105253.

10. Khaledi A, Schniederjans M, Pohl S, Rainer R, Bodenhofer U, Xia B, et al. Transcriptome profiling of antimicrobial resistance in Pseudomonas aeruginosa. Antimicrobial agents and chemotherapy. 2016;60(8):4722–33.

11. O’Rourke A, Beyhan S, Choi Y, Morales P, Chan AP, Espinoza JL, et al. Mechanism-of-action classification of antibiotics by global transcriptome profiling. Antimicrobial agents and chemotherapy. 2020;64(3):10.1128/aac.01207-19.

12. Tatta ER, Paul S, Kumavath R. Transcriptome analysis revealed the synergism of novel rhodethrin inhibition on biofilm architecture, antibiotic resistance and quorum sensing in Enterococcus faecalis. Gene. 2023;871:147436.

13. Liao Z, Lin K, Liao W, Xie Y, Yu G, Shao Y, et al. Transcriptomic analyses reveal the potential antibacterial mechanism of citral against Staphylococcus aureus. Frontiers in microbiology. 2023;14:1171339.

14. Bie L, Zhang M, Wang J, Fang M, Li L, Xu H, et al. Comparative analysis of transcriptomic response of Escherichia coli K-12 MG1655 to nine representative classes of antibiotics. Microbiology Spectrum. 2023;11(2):e00317–23.

15. Mohamad N, Amal MNA, Saad MZ, Yasin ISM, Zulkiply NA, Mustafa M, et al. Virulence-associated genes and antibiotic resistance patterns of Vibrio spp. isolated from cultured marine fishes in Malaysia. BMC veterinary research. 2019;15:1–13.

16. Abd Ghafar SZ, Muthukrishnan S, Zolkeflee NKZ, Natrah I, Abas F. Identification of Metabolites From Halamphora Sp. and Its Correlation With Quorum Sensing Inhibitory Activity via UHPLC-ESI-MS/MS-Based Metabolomics and Molecular Networking. Chemistry & Biodiversity. 2025;22(4):e202402282.

17. Martin M. Cutadapt removes adapter sequences from high-throughput sequencing reads. EMBnet journal. 2011;17(1):10–2.

18. Ewels P, Magnusson M, Lundin S, Käller M. MultiQC: summarize analysis results for multiple tools and samples in a single report. Bioinformatics. 2016;32(19):3047–8.

19. Kopylova E, Noé L, Touzet H. SortMeRNA: fast and accurate filtering of ribosomal RNAs in metatranscriptomic data. Bioinformatics. 2012;28(24):3211–7.

20. Dobin A, Davis CA, Schlesinger F, Drenkow J, Zaleski C, Jha S, et al. STAR: ultrafast universal RNA-seq aligner. Bioinformatics. 2013;29(1):15–21.

21. Liao Y, Smyth GK, Shi W. featureCounts: an efficient general purpose program for assigning sequence reads to genomic features. Bioinformatics. 2014;30(7):923–30.

22. Wu T, Hu E, Xu S, Chen M, Guo P, Dai Z, et al. clusterProfiler 4.0: A universal enrichment tool for interpreting omics data. The innovation. 2021;2(3).

23. Luo W, Brouwer C. Pathview: an R/Bioconductor package for pathway-based data integration and visualization. Bioinformatics. 2013;29(14):1830–1.

24. Bondad-Reantaso MG, Arthur JR, Subasinghe RP. Improving biosecurity through prudent and responsible use of veterinary medicines in aquatic food production. FAO Fisheries and Aquaculture Technical Paper. 2012(547):I.

25. Defoirdt T, Boon N, Bossier P. Can bacteria evolve resistance to quorum sensing disruption? PLoS pathogens. 2010;6(7):e1000989.

26. Azimi S, Klementiev AD, Whiteley M, Diggle SP. Bacterial quorum sensing during infection. Annual review of Microbiology. 2020;74(1):201–19.

27. Chadha J, Harjai K, Chhibber S. Revisiting the virulence hallmarks of Pseudomonas aeruginosa: a chronicle through the perspective of quorum sensing. Environmental microbiology. 2022;24(6):2630–56.

28. Omar NS, Emilia SN, Danish-Daniel M, Iehata S, Ikhsan NFM. Probiotics bacteria as quorum sensing degrader control Aeromonas hydrophila pathogenicity in cultured red hybrid tilapia. Indonesian Aquaculture Journal. 2023;18(1):1–15.

29. Muthukrishnan S, Muthar NI, Zakaria MH, Rukayadi Y, Natrah I. Anti-biofilm and anti-quorum sensing activities of the Red seaweed, Gracilaria changii and its associated bacteria. Journal of Applied Phycology. 2023;35(5):2555–66.

30. Aunins TR, Erickson KE, Chatterjee A. Transcriptome-based design of antisense inhibitors potentiates carbapenem efficacy in CRE Escherichia coli. Proceedings of the National Academy of Sciences. 2020;117(48):30699–709.

31. Asfahl KL, Schuster M. Additive effects of quorum sensing anti-activators on Pseudomonas aeruginosa virulence traits and transcriptome. Frontiers in Microbiology. 2018;8:2654.

32. Kim D, Crippen TL, Jordan HR, Tomberlin JK. Quorum sensing gene regulation in Staphylococcus epidermidis reduces the attraction of Aedes aegypti (L.)(Diptera: Culicidae). Frontiers in microbiology. 2023;14:1208241.

33. Masri A, Khan NA, Zoqratt MZHM, Ayub Q, Anwar A, Rao K, et al. Transcriptome analysis of Escherichia coli K1 after therapy with hesperidin conjugated with silver nanoparticles. BMC microbiology. 2021;21(1):51.

34. Wang J, Chi Z, Zhao K, Wang H, Zhang X, Xu F, et al. A transcriptome analysis of the antibacterial mechanism of flavonoids from Sedum aizoon L. against Shewanella putrefaciens. World Journal of Microbiology and Biotechnology. 2020;36(7):94.

35. Kadosh YS, Muthuraman S, Nisaa K, Ben-Zvi A, Byron DLK, Shagan M, et al. Pseudomonas aeruginosa quorum sensing and biofilm attenuation by a di-hydroxy derivative of piperlongumine (PL-18). Biofilm. 2024;8:100215.

36. Liu L, Zeng X, Zheng J, Zou Y, Qiu S, Dai Y. AHL-mediated quorum sensing to regulate bacterial substance and energy metabolism: A review. Microbiological research. 2022;262:127102.

37. Sabaty M, Adryanczyk Gr, Roustan C, Cuiné S, Lamouroux C, Pignol D. Coproporphyrin excretion and low thiol levels caused by point mutation in the Rhodobacter sphaeroides S-adenosylmethionine synthetase gene. Journal of bacteriology. 2010;192(5):1238–48.

38. Namdari F, Hurtado-Escobar GA, Abed N, Trotereau J, Fardini Y, Giraud E, et al. Deciphering the roles of BamB and its interaction with BamA in outer membrane biogenesis, T3SS expression and virulence in Salmonella. PLoS One. 2012;7(11):e46050.

39. Long S, Su M, Chen X, Hu A, Yu F, Zou Q, et al. Proteomic and Mutant Analysis of Hydrogenase Maturation Protein Gene hypE in Symbiotic Nitrogen Fixation of Mesorhizobium huakuii. International Journal of Molecular Sciences. 2023;24(16):12534.

40. Guo J, Zhu J, Zhao T, Sun Z, Song S, Zhang Y, et al. Survival characteristics and transcriptome profiling reveal the adaptive response of the Brucella melitensis 16M biofilm to osmotic stress. Frontiers in Microbiology. 2022;13:968592.

41. Khazaal S, Al Safadi R, Osman D, Hiron A, Gilot P. Investigation of the polyamine biosynthetic and transport capability of Streptococcus agalactiae: the non-essential PotABCD transporter. Microbiology. 2021;167(12):001124.

42. Zhang C, Li C, Aziz T, Alharbi M, Cui H, Lin L. Screening of E. coli O157: H7 AI-2 QS inhibitors and their inhibitory effect on biofilm formation in combination with disinfectants. Food Bioscience. 2024;58:103821.

43. Poole K. Bacterial stress responses as determinants of antimicrobial resistance. Journal of Antimicrobial Chemotherapy. 2012;67(9):2069–89.

44. Baharoglu Z, Mazel D. SOS, the formidable strategy of bacteria against aggressions. FEMS microbiology reviews. 2014;38(6):1126–45.

45. Butler MT, Wang Q, Harshey RM. Cell density and mobility protect swarming bacteria against antibiotics. Proceedings of the National Academy of Sciences. 2010;107(8):3776–81.

46. Porter SL, Wadhams GH, Armitage JP. Signal processing in complex chemotaxis pathways. Nature Reviews Microbiology. 2011;9(3):153–65.

47. Giannakara M, Koumandou VL. Evolution of two-component quorum sensing systems. Access microbiology. 2022;4(1):000303.

48. Sionov RV, Steinberg D. Targeting the holy triangle of quorum sensing, biofilm formation, and antibiotic resistance in pathogenic bacteria. Microorganisms. 2022;10(6):1239.

49. Alvarez AF, Georgellis D. Environmental adaptation and diversification of bacterial two-component systems. Current Opinion in Microbiology. 2023;76:102399.

50. Wang L, Pan Y, Yuan Z-H, Zhang H, Peng B-Y, Wang F-F, et al. Two-component signaling system VgrRS directly senses extracytoplasmic and intracellular iron to control bacterial adaptation under iron depleted stress. PLoS pathogens. 2016;12(12):e1006133.

51. Beier D, Gross R. Regulation of bacterial virulence by two-component systems. Current opinion in microbiology. 2006;9(2):143–52.

52. Phoka T, Fule L, Da Fonseca JP, Cokelaer T, Picardeau M, Patarakul K. Investigating the role of the carbon storage regulator A (CsrA) in Leptospira spp. Plos one. 2021;16(12):e0260981.

53. Dai Q, Xu L, Xiao L, Zhu K, Song Y, Li C, et al. RovM and CsrA negatively regulate urease expression in Yersinia pseudotuberculosis. Frontiers in Microbiology. 2018;9:348.

54. Gu H, Qi H, Chen S, Shi K, Wang H, Wang J. Carbon storage regulator CsrA plays important roles in multiple virulence-associated processes of Clostridium difficile. Microbial pathogenesis. 2018;121:303–9.

55. Zhang Z, Zhao Y, Chen X, Li W, Li W, Du J, et al. Effects of cinnamon essential oil on oxidative damage and outer membrane protein genes of Salmonella enteritidis cells. Foods. 2022;11(15):2234.

56. Aung HL, Berney M, Cook GM. Hypoxia-activated cytochrome bd expression in Mycobacterium smegmatis is cyclic AMP receptor protein dependent. Journal of bacteriology. 2014;196(17):3091–7.

57. Ko E-M, Oh J-I. Induction of the cydAB operon encoding the bd quinol oxidase under respiration-inhibitory conditions by the major cAMP receptor protein MSMEG_6189 in Mycobacterium smegmatis. Frontiers in microbiology. 2020;11:608624.

58. Poole K. Stress responses as determinants of antimicrobial resistance in Gram-negative bacteria. Trends in microbiology. 2012;20(5):227–34.

59. Claverys J-P, Prudhomme M, Martin B. Induction of competence regulons as a general response to stress in gram-positive bacteria. Annu Rev Microbiol. 2006;60(1):451–75.

60. Hancock V, Klemm P. Global gene expression profiling of asymptomatic bacteriuria Escherichia coli during biofilm growth in human urine. Infection and immunity. 2007;75(2):966–76.

61. Du Plessis DJ, Nouwen N, Driessen AJ. The sec translocase. Biochimica et Biophysica Acta (BBA)-Biomembranes. 2011;1808(3):851–65.

62. Liang W, Pascual-Montano A, Silva AJ, Benitez JA. The cyclic AMP receptor protein modulates quorum sensing, motility and multiple genes that affect intestinal colonization in Vibrio cholerae. Microbiology. 2007;153(9):2964–75.

63. Zheng D, Constantinidou C, Hobman JL, Minchin SD. Identification of the CRP regulon using in vitro and in vivo transcriptional profiling. Nucleic acids research. 2004;32(19):5874–93.

64. Martínez JL, Rojo F. Metabolic regulation of antibiotic resistance. FEMS microbiology reviews. 2011;35(5):768–89.

65. Zhang YJ, Reddy MC, Ioerger TR, Rothchild AC, Dartois V, Schuster BM, et al. Tryptophan biosynthesis protects mycobacteria from CD4 T-cell-mediated killing. Cell. 2013;155(6):1296–308.

66. Rutherford ST, Van Kessel JC, Shao Y, Bassler BL. AphA and LuxR/HapR reciprocally control quorum sensing in vibrios. Genes & development. 2011;25(4):397–408.

67. Deng Y, Chen C, Zhao Z, Zhao J, Jacq A, Huang X, et al. The RNA chaperone Hfq is involved in colony morphology, nutrient utilization and oxidative and envelope stress response in Vibrio alginolyticus. PLoS One. 2016;11(9):e0163689.

68. Vogel J, Luisi BF. Hfq and its constellation of RNA. Nature Reviews Microbiology. 2011;9(8):578–89.

69. Teplitski M, Goodier RI, Ahmer BM. Pathways Leading from BarA/SirA to Motility andVirulence Gene Expression in Salmonella. Journal of bacteriology. 2003;185(24):7257–65.

70. Wang Q, Frye JG, McClelland M, Harshey RM. Gene expression patterns during swarming in Salmonella typhimurium: genes specific to surface growth and putative new motility and pathogenicity genes. Molecular microbiology. 2004;52(1):169–87.

71. Wei BL, Brun-Zinkernagel AM, Simecka JW, Prüß BM, Babitzke P, Romeo T. Positive regulation of motility and flhDC expression by the RNA-binding protein CsrA of Escherichia coli. Molecular microbiology. 2001;40(1):245–56.

72. Postma PW, Lengeler JW, Jacobson G. Phosphoenolpyruvate: carbohydrate phosphotransferase systems of bacteria. Microbiological reviews. 1993;57(3):543–94.

73. Provenzano D, Klose KE. Altered expression of the ToxR-regulated porins OmpU and OmpT diminishes Vibrio cholerae bile resistance, virulence factor expression, and intestinal colonization. Proceedings of the National Academy of Sciences. 2000;97(18):10220–4.

74. Schneider R, Travers A, Kutateladze T, Muskhelishvili G. A DNA architectural protein couples cellular physiology and DNA topology in Escherichia coli. Molecular microbiology. 1999;34(5):953–64.

75. Battesti A, Majdalani N, Gottesman S. The RpoS-mediated general stress response in Escherichia coli. Annual review of microbiology. 2011;65(1):189–213.

76. Yildiz FH, Schoolnik GK. Vibrio cholerae O1 El Tor: identification of a gene cluster required for the rugose colony type, exopolysaccharide production, chlorine resistance, and biofilm formation. Proceedings of the National Academy of Sciences. 1999;96(7):4028–33.

